# Body condition score and triglyceride concentrations and their associations with other markers of energy homeostasis in healthy, non-obese dogs

**DOI:** 10.1101/2022.09.19.508523

**Authors:** Carlos Gomez-Fernandez-Blanco, Dominique Peeters, Frédéric Farnir, Katja Höglund, Vassiliki Gouni, Maria Wiberg, Jakob Lundgren Willesen, Sofia Hanås, Kathleen McEntee, Laurent Tiret, Jens Häggström, Hannes Lohi, Valérie Chetboul, Merete Fredholm, Eija Seppälä, Anne-Sophie Lequarré, Alexander James German, Anne-Christine Merveille

## Abstract

Serum triglyceride concentrations increase in dogs in overweight condition, which is typically assessed by body condition score (BCS). However, their associations with other markers of energy homeostasis are poorly characterized. The present study aimed to evaluate the associations between both BCS and triglyceride levels and other markers of lipid and glucose metabolism in healthy dogs in overweight condition. 534 overweight, but otherwise healthy, client-owned dogs were included. Serum concentrations of cholesterol, free fatty acids, triglycerides, insulin, glucose and fructosamine were measured. Dogs were assigned to lean (BCS: 3-5) or overweight (BCS: 6-7) categories, and linear models were used to assess the differences between BCS categories and the associations between triglycerides and the other variables, correcting for the effect of breed. Globally, “overweight” dogs had greater serum cholesterol (95% CI: 5.3-6.2 mmol/L or 205-237 mg/dL versus 5.1-5.4 mmol/L or 198-210 mg/dl, P = .003), insulin (95% CI: 17.5-22.1 μU/ml versus 16.7-18.0 μU/ml, P = .036) and were older (95% CI: 4.0-5.3 versus 3.4-3.7 years, P = .002) than lean dogs. Triglyceride concentrations were positively associated with fructosamine (r2 = 0.31, P = .001), cholesterol (r2 = 0.25, P < .001), insulin (r2 = 0.14, P = .003) and glucose (r2 = 0.10, P = .002), and negatively associated with free fatty acids (r2 = 0.11, P < .001). There was no association between triglyceride levels and age. In conclusion, both BCS and triglyceride concentrations were associated with other markers of glucose and lipid metabolism in overweight, but otherwise healthy dogs. Triglyceride concentrations were associated with an increase in insulin and fructosamine that might reflect an early-phase impairment in glucose tolerance which, surprisingly, was concurrent with lower basal free fatty acids.

## Introduction

Metabolic syndrome (MS) is an entity comprising multiple cardiovascular and metabolic risk factors in humans, characterized by a state of chronic, subclinical inflammation (1). Features of human MS include combinations of increased visceral fat (abdominal obesity), systemic hypertension, increased circulating triglyceride (TG) concentrations, decreased high-density lipoprotein (HDL) cholesterol concentration, and increased fasting glucose concentration (suggesting insulin resistance) (2). Human beings with MS have a greater risk of developing type 2 diabetes mellitus (DM) and cardiovascular complications (2).

Dogs also suffer from obesity-related metabolic dysfunction (ORMD), but the term metabolic syndrome is avoided because, even though dogs share some of its components (such as hypercholesterolaemia, hypertriglyceridaemia and insulin resistance), they do not develop the obesity-related diseases that humans do, such as atherosclerosis, stroke or type 2 DM (3–6). Studies on canine ORMD have mostly examined dogs with manifest obesity, whilst metabolic dysfunction of dogs in overweight condition has been less well characterized, despite its known associated with comorbidities in the canine species (7–10).

Some people with obesity remain metabolically healthy, at least in the short term (11), during which time their risk of comorbidities is less, they do not develop insulin resistance, are normotensive, and concentration of glucose, TG, HDL cholesterol and high-sensitivity C-reactive protein (hsCRP) is within reference limits (12). Conversely, some normal-weight individuals develop the characteristics of MS despite having a normal body fat mass (13). Such ‘metabolically unhealthy normal-weight’ people can be identified by a thorough biochemical assessment in which increased TG and C-reactive protein (CRP) concentration, as well as decreased HDL-cholesterol and adiponectin concentrations will be found (13). Similarly, different biochemical phenotypes have been described in obese dogs: adiponectin was lesser and plasma insulin was greater in obese dogs that met the criteria of ORMD (which were defined as having obesity plus two other criteria amongst TG > 200 mg/dL (2.26 mmol/L), total cholesterol > 300mg/dL (7.8 mmol/L), systolic blood pressure >160 mmHg, and fasting plasma glucose > 100mg/dL (5.6 mmol/L), or previously diagnosed diabetes mellitus) as compared to obese dogs without ORMD (14,15).

The aim of this study was to investigate associations amongst body condition score (BCS) and a range of metabolic variables associated with glucose and fat metabolism in a large cohort of healthy dogs in overweight condition, as compared to lean dogs, as well as to assess whether TG concentrations can be a marker for these metabolic variables.

## Materials and methods

A canine database aimed at examining genetic determinants of disease (European LUPA project (16)) was retrieved. Five centres had participated in the original study between 2009 and 2010: University of Liège, University of Copenhagen, Swedish University of Agricultural Sciewnces, University of Helsinki, and the National Veterinary School of Maisons-Alfort (France). All centres used the same standardised protocols for recruitment and characterisation of dogs. The database was then retrospectively investigated for the purpose of the present work.

### Dogs

This study was approved by the Ethical Committee of the LUPA project, and the informed consent of all owners was obtained (16). Client-owned, pure-bred dogs were recruited, and included different breed cohorts in order to represent a wide range of various phenotypic features. Dogs were 1 to 7 years old and were genetically unrelated. In order to minimise potential effects of the oestrous cycle on metabolic parameters, each breed cohort comprised a single sex, namely intact males or female dogs that were spayed or in anoestrus (checked by a vaginal smear). Health status was checked through history, physical examination, laboratory analyses (including routine hematology and serum biochemistry), and a thorough cardiovascular investigation comprising ECG recording and echocardiographic examination. After visual assessment and palpation, each dog was assigned a BCS category using the one-to-nine point scale (17); dogs were then assigned to one of two body condition categories (lean, BCS 3-5; overweight, BCS 6-7) based on this score. Dogs were excluded if they showed clinical signs of any disease, were very underweight (BCS ≤2) or had obesity (BCS 8-9, as defined by Brooks et al. (18)).

### Sampling

Three weeks before the study, owners were asked to feed their dogs exclusively with a commercial dry food diet of their choice, avoiding any treats or table food. Dogs were fasted for at least 12 hours before blood sampling and, after collection, blood was centrifuged within 30 minutes and serum then aliquoted. In most centres, serum aliquots were immediately frozen at −80°C; in the remaining centre, samples were frozen at −20°C for the first 2 weeks after collection, before being transferred to a −80°C thereafter. All samples were subsequently sent to the same laboratory^a^ for analysis.

### Analyses

The analytes selected for the present study were chosen based on their reported association with obesity in humans and dogs (2,3,14,19–41). These included markers of lipid metabolism (e.g., cholesterol, TG and free fatty acids, FFA), glucose homeostasis (e.g., glucose, fructosamine, and insulin) and inflammation (CRP). A photometric clinical chemistry analyzer (Konelab 60i, Thermo Electron Co, Finland) was used to determine fructosamine, glucose, CRP, cholesterol and TG concentrations. Fructosamine and FFA concentrations were respectively assessed using Hariba ABX (Montpellier, France) and Wako Chemical Gmbh (Neuss, Germany). Insulin concentration was determined by radioimmunoassay, provided by DiaSorin S.p.A (Italy).

### Repeatability of measurements

Ten dogs were randomly selected to send duplicates to the same laboratory three months after the first analyses. Coefficients of variation were 6% for insulin and ≤5% for FFA, cholesterol, TG, glucose and fructosamine.

### Statistical analyses

Outliers were inspected with the Reference Value Advisor (42) using the Tukey method (43). Dogs with only one outlier were accepted and included in the analyses; those with more than one outlying value were excluded.

### Preliminary tests

Given that parametric tests were needed to test our hypotheses, all analyses were validated after checking the normal distribution of the residuals (evaluated by histogram observation and by a Shapiro-Wilk test), and a test of homoscedaticity (Breusch-Pagan test). If a particular analyte (dependent variable) did not pass the normality tests, a non-parametric equivalent test was used (e.g., Spearman’s Rank correlation or Mann-Whitney test). Statistical tests were performed with R free statistical software (44).

### Effect of age on biochemical variables

The correlation between age and each of the analytes was calculated by Pearson’s method, and *P*-values <0.05 were considered to be statistically significant.

### Effect of overweight status

In order to test for differences between lean and overweight groups, a linear model including BCS, breed and an interaction between BCS and breed was used for all biochemical variables. Normality was checked using the Shapiro-Wilk test and analytes with non-normal residuals were log-transformed. Type III sums of squares were used and differences with *P*-values <0.05 were considered to be statistically significant. Whenever a statistically significant interaction was found, associations within each breed were examined. In this case, a Mann-Whitney test was used and the level of significance was corrected for multiple comparisons using a Bonferroni correction.

### Effect of TG on the biochemical variables

Linear models including TG, the breed and an interaction between the breed and TG were used for each biochemical variable. Once again, *P*-values <0.05 were considered to be statistically significant. Whenever an interaction with the breed was significant, the association between TG and the outcome variable was tested within each breed, using Spearman’s correlation method. Once again, a Bonferroni correction was used to correct for multiple comparisons.

## Results

### Animals

In total, 534 dogs met the inclusion criteria, with 9 different breeds represented: Boxer (BOX, 15 dogs), Belgian Shepherd Dog (BSD, 125 dogs), Cavalier King Charles Spaniel (CKCS, 35 dogs), Dachshund (DACH, 40 dogs), Doberman (DOB, 39 dogs), Finnish Lapphund (FL, 45 dogs), German Shepherd dog (GSD, 66 dogs), Labrador Retriever (LAB, 125 dogs), and Newfoundland (NF, 44 dogs). All cohorts comprised only male dogs, except for the NF cohort, that comprised only females, and the LAB cohort, that comprised dogs of both sexes (73 females and 52 males). Some breeds were unique to one centre while others were shared among centres. Distribution of dogs by centre, breed, and gender is shown in Table 1. All outliers were used in the statistical analyses, since they were considered compatible with physiological values, and no dog showed more than one outlier result.

**Table 1.**
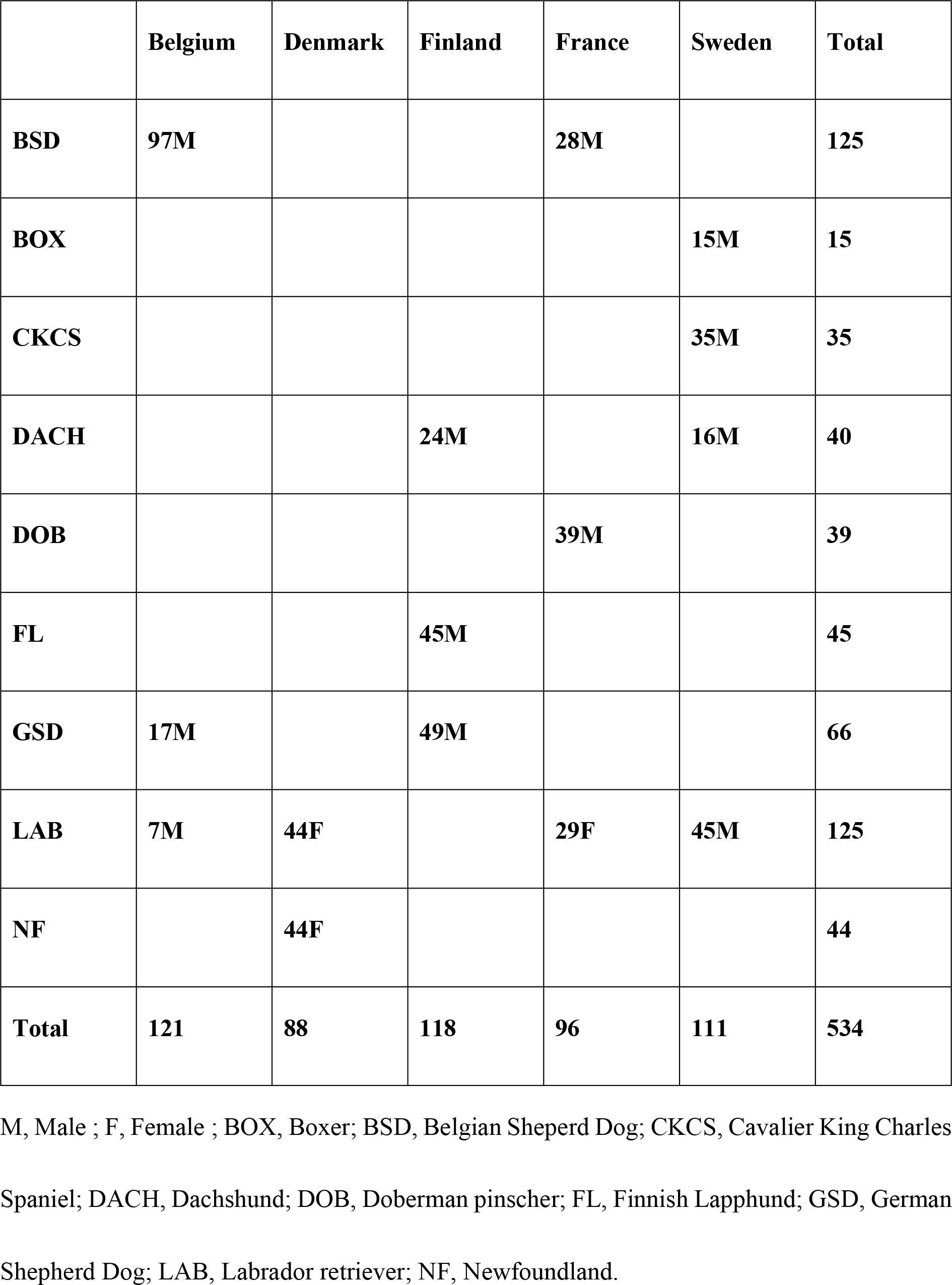
Distribution of dogs by centre, breed, and sex

Median BCS was 3 in FL (interquartile range, IQR: 3-4); 4 in GSD (IQR: 3-4), DACH (IQR: 3-5), BSD (IQR: 4-5) and BOX (IQR: 4-5); 5 in DOB (IQR: 5-5), CKCS (IQR: 5-6) and NF (IQR: 5-6); and 5.5 in LAB (IQR: 5-6). Seventy-one percent of the dogs (409) were in the lean category, while 29% (120 dogs) were overweight. Table 2 shows the distribution of BCS amongst breeds.

**Table 2.** Distribution of dogs by breed and BCS.

|  | **BCS 2** | **BCS 3** | **BCS 4** | **BCS 5** | **BCS 6** | **BCS 7** |
| --- | --- | --- | --- | --- | --- | --- |
| **BOX** | - | - | 8 | 4 | 3 | - |
| **BSD** | 1 | 22 | 45 | 51 | 5 | 1 |
| **CKCS** | - | - | - | 20 | 15 | - |
| **DACH** | - | 11 | 12 | 11 | 6 | - |
| **DOB** | - | - | 2 | 33 | 4 | - |
| **FL** | - | 24 | 12 | 6 | 2 | 1 |
| **GS** | 1 | 22 | 35 | 4 | 1 | - |
| **LAB** | - | 1 | 8 | 53 | 59 | 4 |
| **NF** | - | - | - | 23 | 15 | 4 |
| **TOTAL** | **2** | **80** | **122** | **205** | **110** | **10** |

BCS, Body Condition Score ; BOX, Boxer; BSD, Belgian Sheperd Dog; CKCS, Cavalier King Charles Spaniel; DACH, Dachshund; DOB, Doberman pinscher; FL, Finnish Lapphund; GSD, German Shepherd Dog; LAB, Labrador retriever; NF, Newfoundland.

Analyses were validated after successfully testing for normal distribution of the residuals and homoscedaticity. CRP was the only variable that did not succeed the tests and was, therefore, assessed with nonparametric methods.

### Association between age and biochemical variables

There was no effect of age on any analyte tested (cholesterol, *P*=0.49; FFA, *P*=0.88; TG, *P*=0.55; CRP, *P*=0.34; insulin, *P*=0.18; glucose, *P*=0.65; fructosamine, *P*=0.11).

### Effect of overweight status on biochemical variables

After normalisation for the effect of the breed, dogs in the overweight category were older (*P*=0.002) and had greater plasma insulin (*P*=0.036) and cholesterol (*P*=0.003) concentrations than lean dogs. An interaction between BCS category and breed was also identified for both TG and cholesterol (Table 3): in this respect, cholesterol concentration was greater in overweight BOX (*P*=0.020) and CKCS (*P*=.0005) compared with their lean counterparts; overweight CKCS also had greater TG than lean CKCS (*P*=0.002). These differences are illustrated in Fig 1.

**Table 3.** Results of ANOVA. Effect of BCS category, and its interaction with the effect of the breed, on age (years) and on concentrations of insulin, fructosamine, glucose, cholesterol, free fatty acids, and triglycerides in overweight and lean dogs. Data are presented as mean and 95% confidence intervals.

|  | Lean | Overweight | BCS<br>category, p-value | Interaction<br>between BCS<br>category and<br>breed, p- value |
| --- | --- | --- | --- | --- |
| Age (years) | 3.5<br>(3.4-3.7) | 4.4<br>(3.9-4.9) | 0.0005<sup>a</sup> | 0.3405 |
| Insulin (pmol/l) | 120.1<br>(113.2-125.7) | 131.3<br>(122.2-141.7) | 0.0374<sup>a</sup> | 0.1289 |
| **Insulin (μU/ml)** | 17.3<br>(16.7-18.1) | 18.9<br>(17.6-20.4) |  |  |
| **Fructosamine (μmol/l)** | 287.1<br>(283.6 – 290.6) | 289.7<br>(283.2 – 296.2) | 0.4951 | 0.4560 |
| **Glucose (mmol/l)** | 0.05<br>(0.05-0.05) | 0.05<br>(0.05-0.06) | 0.3828 | 0.6783 |
| **Glucose (mg/dl)** | 0.98<br>(0.97-0.99) | 0.99<br>(0.97-1.01) |  |  |
| **Cholesterol (mmol/l)** | 5.3<br>(5.1-5.4) | 5.7<br>(5.3-6.2) | 0.0032<sup>a</sup> | 0.0006<sup>b</sup> |
| **Cholesterol (mg/dl)** | 204<br>(198-210) | 221<br>(205-237) |  |  |
| **FFA (mEq/l)** | 0.87<br>(0.83-0.91) | 0.86<br>(0.78-0.93) | 0.7657 | 0.7529 |
| **Triglycerides**<br>**(mmol/l)** | 0.005<br>(0.005-0.005) | 0.005<br>(0.005-0.006) | 0.9716 | 0.0019<sup>b</sup> |
| **Triglycerides**<br>**(mg/dl)** | 0.43<br>(0.42-0.45) | 0.45<br>(0.40-0.51) |  |  |

FFA, free fatty acids.
<sup>a</sup> Significant effect of the BCS category for a level of confidence of 0.05.
<sup>b</sup> Significant interaction between breed and BCS category, for a level of confidence of 0.05.

**Fig. 1.**
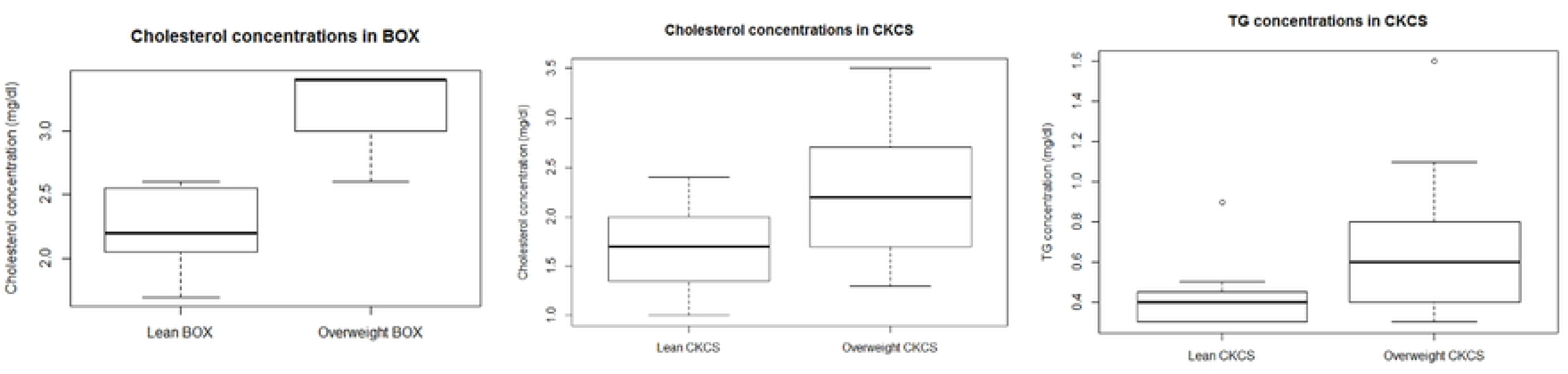
Box plots showing serum cholesterol (A and B) and triglyceride (C) concentrations in Boxer (BOX) and Cavalier King Charles Spaniel (CKCS). Bonferroni-corrected p-value of 0.0056. The lower, middle and upper line of each box represent the 25^th^ percentile (bottom quartile), 50^th^ percentile (median) and the 75^th^ percentile (top quartile). The whiskers, where present, represent the minimum and maximum. Outliers, represented by open circles, were included in the analyses.

### Associations between TG concentration and markers of ORMD

After normalisation for the effect of the breed, TG concentrations were positively associated with fructosamine (*P*=0.001), cholesterol *(P*<0.001), insulin (*P*=0.003) and glucose (*P*=0.001) concentrations, and negatively associated with FFA (*P*<.0001). Results are shown in **Table 4**. The interaction between TG and breed was significant when testing insulin concentration as the dependent variable and, in the intra-breed analyses, there was a positive association between TG and insulin concentrations in several breeds (**Table 5**).

**Table 4.**
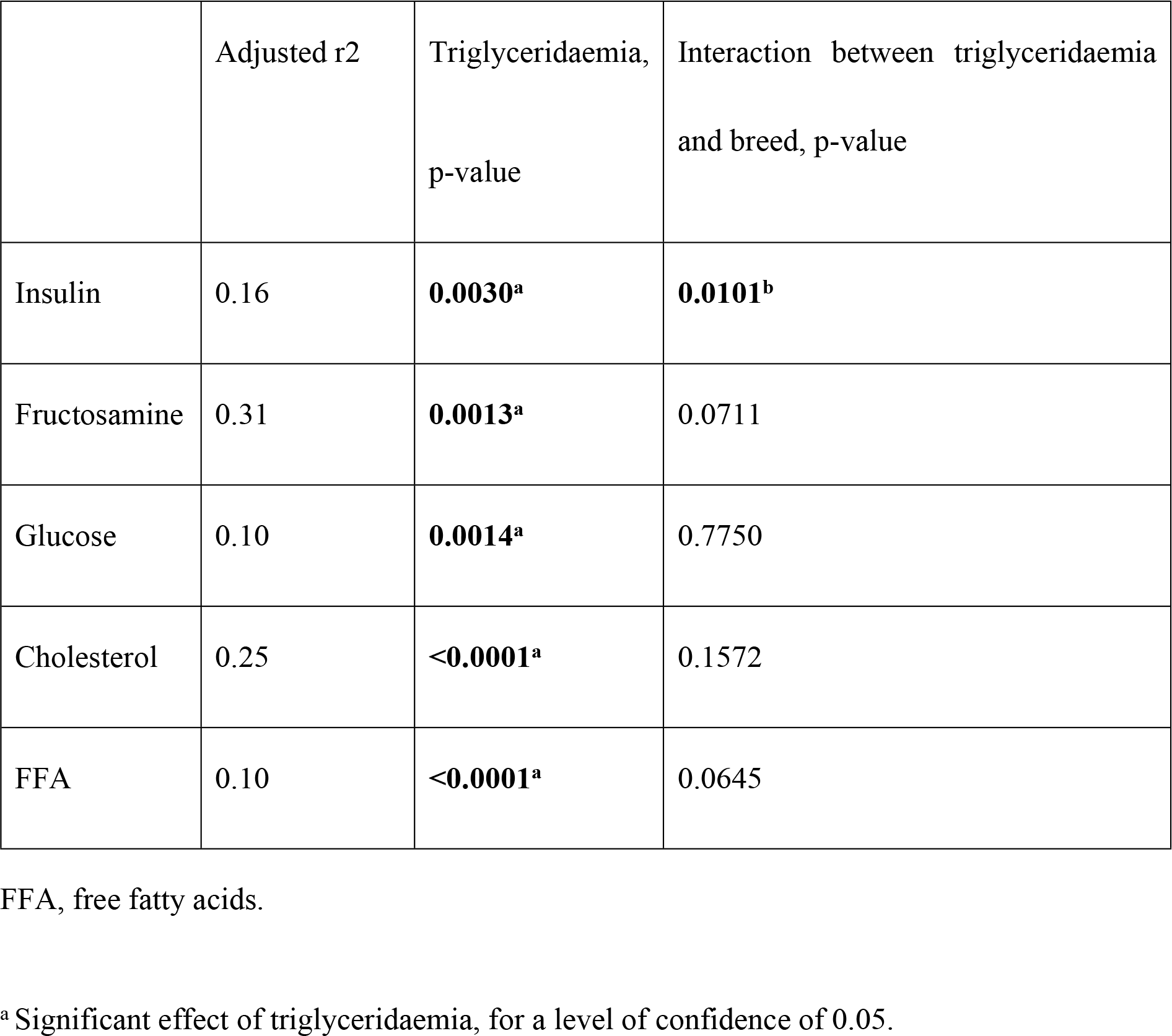
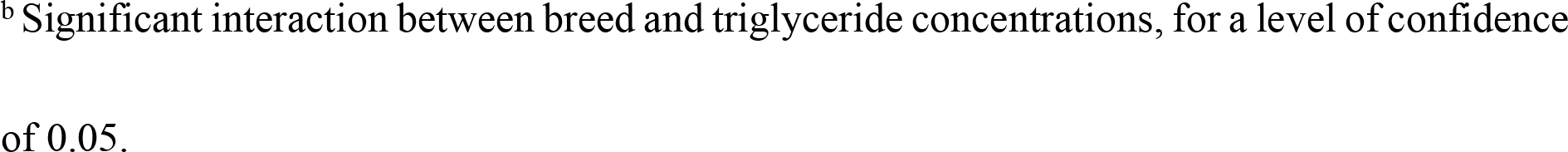
Association between triglyceride concentration and insulin, fructosamine, glucose, cholesterol, and free fatty acids. The r^2^ corresponds to the predictive percentage attributed to the whole linear model, which included the main effects of both triglycerides and breed (plus their interaction, when significant) as explanatory variables.

| | Adjusted $r^2$ | Triglyceridaemia,<br>p-value | Interaction between triglyceridaemia<br>and breed, p-value |
| --- | --- | --- | --- |
| Insulin | 0.16 | **0.0030<sup>a</sup>** | **0.0101<sup>b</sup>** |
| Fructosamine | 0.31 | **0.0013<sup>a</sup>** | 0.0711 |
| Glucose | 0.10 | **0.0014<sup>a</sup>** | 0.7750 |
| Cholesterol | 0.25 | **<0.0001<sup>a</sup>** | 0.1572 |
| FFA | 0.10 | **<0.0001<sup>a</sup>** | 0.0645 |

FFA, free fatty acids.
<sup>a</sup> Significant effect of triglyceridaemia, for a level of confidence of 0.05.
<sup>b</sup> Significant interaction between breed and triglyceride concentrations, for a level of confidence of 0.05.

**Table 5.**
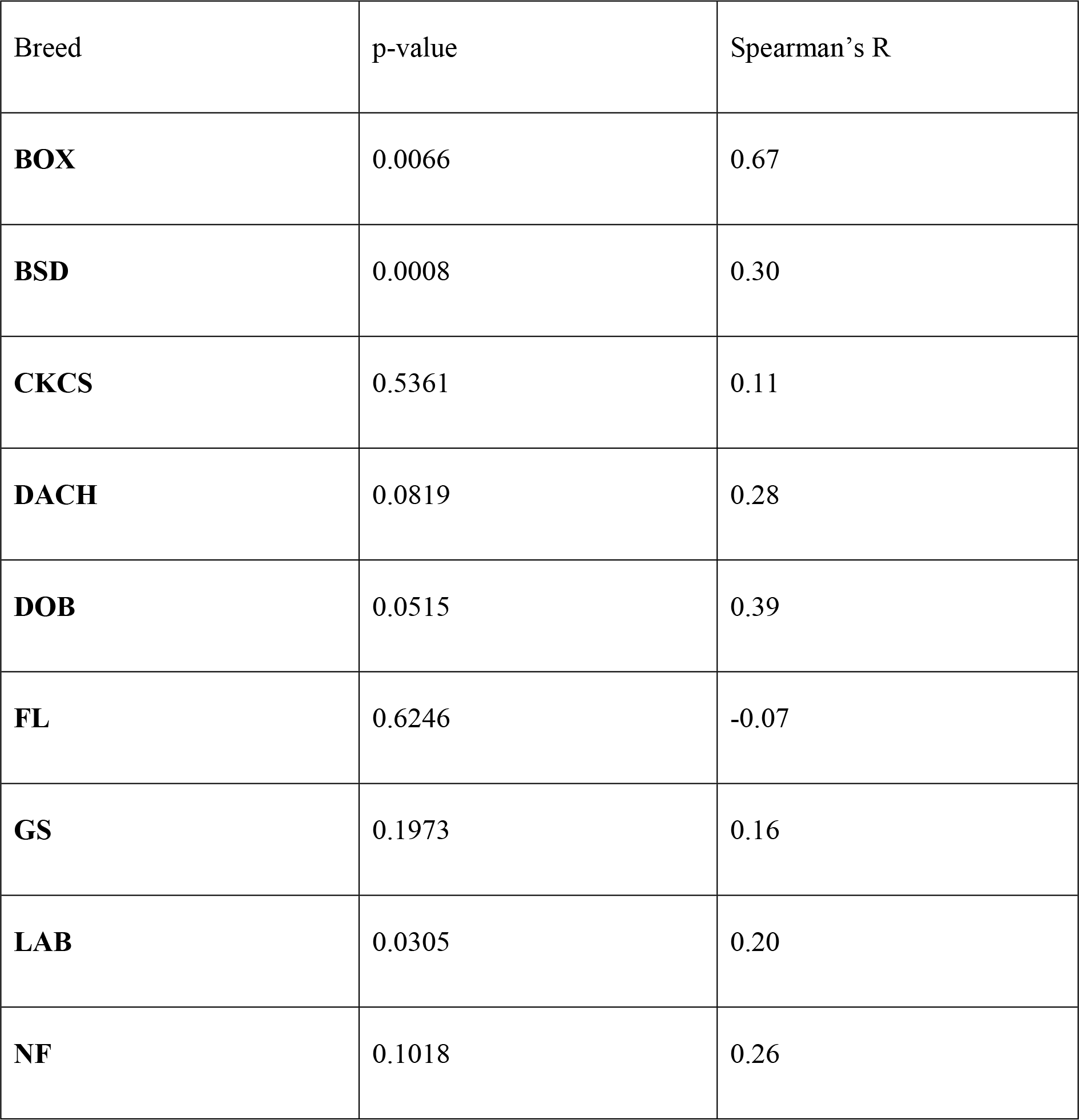
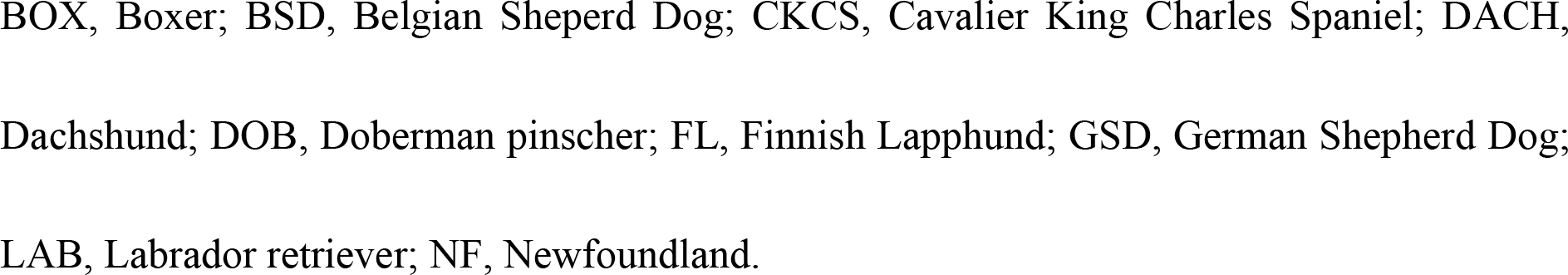
Correlations between triglyceride concentration (mg/dl) and insulin (μU/ml) within individual breeds. Spearman’s rank correlation test.

| Breed | p-value | Spearman's R |
| --- | --- | --- |
| **BOX** | 0.0066 | 0.67 |
| **BSD** | 0.0008 | 0.30 |
| **CKCS** | 0.5361 | 0.11 |
| **DACH** | 0.0819 | 0.28 |
| **DOB** | 0.0515 | 0.39 |
| **FL** | 0.6246 | -0.07 |
| **GS** | 0.1973 | 0.16 |
| **LAB** | 0.0305 | 0.20 |
| **NF** | 0.1018 | 0.26 |

BOX, Boxer; BSD, Belgian Sheperd Dog; CKCS, Cavalier King Charles Spaniel; DACH, Dachshund; DOB, Doberman pinscher; FL, Finnish Lapphund; GSD, German Shepherd Dog; LAB, Labrador retriever; NF, Newfoundland.

## Discussion

Two main findings can be highlighted from this large cohort of dogs recruited from various European countries. First, insulin and cholesterol concentrations are increased in dogs in overweight body condition; second, fasting TG concentrations are positively associated with cholesterol, glucose, fructosamine and insulin, but negatively associated with FFA. Previous studies, often experimental, have reported increased insulin, cholesterol and TG concentrations in obese dogs. However, the “overweight” group in those studies invariably comprises either some or all dogs in obese body condition (BCS 8-9) (14,19,23,35–38,45–47). To the best of our knowledge, the present study is the first to report similar changes in a large group of dogs in overweight condition only (BCS 6-7). Currently, consensus does not exist on when accumulated adipose tissue becomes pathologicial in dogs and, in some previous studies, dogs with BCS 6/9 have been included in the “ideal weight” group, together with BCS 4 and BCS 5 (32,48,49). Conversely, some studies have examined “mildly to moderately overweight” dogs separately from “obese” dogs and found both groups to be at greater risk for developing comorbidities, suffering from a poorer quality of life and having a shorter life-expectancy (7–10). For example, Kealy et al. (2002) found that long-term food-restricted Labrador Retriever dogs had a longer life span and delayed onset of chronic diseases as compared to a control group (8). There was a mean difference of 26% between groups, which was reflected by a difference in BCS (mean BCS 4.6 +/− 0.19 in the food-restricted group, versus BCS 6.7 +/− 0.19 in the control group). In the present study, insulin and cholesterol concentrations were significantly greater in a cohort of overweight dogs (median = 6; IQR = 0; mean BCS = 6.1; range, 6 – 7) than in their lean counterparts (median BCS = 5; IQR =1; mean BCS = 4.3; range, 2 – 5).

In addition to the main effects, both TG and cholesterol concentrations were affected by an interaction between overweight status and breed. When such an interaction between two independent variables is found, interpretation of the main effects alone may be misleading and, therefore, each category (in this case, individual breeds) should be investigated independently (50). In this respect, compared with lean status, overweight status was associated with greater TG concentrations in the CKCS, and also greater cholesterol concentrations in both BOX and CKCS breeds. We hypothesize that the positive results within these breeds might be related to the experimental design of the present study, rather than a breed-specific cause. Hyperinsulinaemia is thought to be key in obesity-related disorders in humans (51). Even though hyperinsulinaemia is also a feature of canine obesity, dogs do not develop the same outcomes as humans with MS (14), suggesting that significant physiological and pathophysiological differences might exist between these species (3). In the current study, overweight status in dogs was associated with increased concentrations of insulin and cholesterol, which might be interpreted as an early evidence of ORMD. These changes, although mild and likely not clinically relevant, might contribute to the long-term consequences of fat accumulation (i.e., reduced lifespan and quality of life, rapid onset of comorbidities) that have been described in overweight dogs (7–10).

In contrast to TG and cholesterol, FFA concentrations were not different between dogs in the overweight and lean categories. Previous studies have shown that both humans and dogs with obesity have increased FFA concentrations, possibly because their concurrent insulin resistance leads to a lack of insulin-mediated suppression of lipolysis (36,37,46,51); further, FFA are considered to be key mediators in the pathogenesis of obesity-induced insulin resistance (51). Therefore, the degree of adiposity in the dogs of the current study might have been less than that required to affect circulating FFA concentrations.

Fasting hyperglycaemia is seen in humans with obesity (52), and is a risk factor for progression to type 2 DM (2). In dogs, type 2 DM has not been convincingly described (53,54), but the incidence of diabetes mellitus has increased in the last years, paralleling the increase in obesity (8,55,56), whilst several studies report obesity to be a risk factor (57,58).

In the current study, there was no difference in glucose concentrations between the dogs in the overweight and lean categories. Several canine studies have identified changes in glucose concentration associated with overweight status, weight gain and weight loss (39–41,59), but no such association is evident in other studies (14,19–21,24,35,37,60,61). In the present study, fructosamine concentration was did not differ between the dogs in the overweight and lean categories.

Fructosamine is not often included in studies on canine obesity. One study found greater fructosamine concentrations in insulin-resistant, but not insulin-sensitive, dogs with obesity (27). The influence of obesity on fructosamine concentrations in humans is believed to be mild (28,29).

Some of the breeds included in the current study were overrepresented in the overweight category. This finding was expected for both LAB (62,63) and NF (18), breeds that have previously been reported to be predisposed to obesity (18,30,62,63). In contrast, some other obesity-prone breeds were under-represented in the overweight category, for example, there were only 15% and 20% of overweight dogs in the DACH and BOX categories, respectively (30).

There was no difference in CRP concentration between dogs in the overweight and lean categories. CRP is a marker of inflammation and is increased in humans with obesity (31). Subclinical inflammation is also a feature of obesity in dogs, and some authors have found a positive association between CRP and either obesity or weight loss in dogs (20,32). However, this finding is inconsistent, and it was not evident in other studies (33,35,41,64). In one study, CRP concentrations were less in dogs with obesity (34).

When assessing TG as an independent marker of ORMD in the dogs of the current study, positive associations with glucose, fructosamine, insulin and cholesterol were identified, whilst TG concentration was negatively associated with FFA. In people, plasma TG concentration is an independent predictor of MS (65). TG concentration, both independently and alongside other biochemical variables (such as insulin, hsCRP, adiponectin and HDL-cholesterol), can predict a greater risk of type 2 DM and cardiovascular complications even in normal-weight humans, and this analyte might be a more sensitive marker than the common definition of MS (66). Also, studies in children have shown that a decrease in TG concentration following a low-fat diet, was associated with healthier metabolic profiles even though no significant changes in BCS were observed (67).

In one study, dogs with concurrent obesity were grouped according to the presence of metabolic dysfunction using criteria similar to those of human MS (14); dogs were classified as having so-called ORMD when BCS was between 7 and 9 and there were abnormalities in at least two of the following: triglycerides, total cholesterol, systolic blood pressure and fasting plasma glucose concentrations (14). The dogs classified in this way as ORMD did not have a greater total fat mass than those without ORMD, but insulin concentration was greater and adiponectin concentration was lower than in obese dogs not meeting the criteria for ORMD. This suggested that the assessment of metabolic risk could help to classify dogs at risk of obesity-associated comorbidities. According to the results of the current study, plasma TG concentration, used as an only explanatory variable and correcting for the effect of the breed, were associated with metabolic variables but not BCS, including greater glucose and fructosamine concentrations. Of course, this finding should be interpreted in light of the fact that none of the dogs were in obese body condition (BCS 8 to 9). To the author’s knowledge, an association between TG and glucose and fructosamine has not been previously reported in dogs, and might suggest an association between increased glucose concentration and altered lipid metabolism. Fructosamine is seldom researched in studies of human MS, but has been linked to cardiovascular outcomes, whilst its concentration increases with dyslipidaemia, including an association with triglyceride concentration (68).

Another important finding of the present study was the association between fasting TG and insulin concentrations which, to the best of the authors’ knowledge, has not previously been described in healthy dogs. Increased TG concentrations are commonly associated with insulin resistance and type 2 DM in humans, and are considered to be the central feature of the dyslipidaemia that is present in these states (69–71). In dogs, it has been suggested that hypertriglyceridaemia is favoured by an increased supply of substrates to the liver (especially glucose and FFA) in insulin-resistant states (36).

In a study involving dogs with obesity, insulin concentration was less in a subgroup of persistently-hyperglycaemic dogs compared with obese dogs that were not persistently hyperglyacemic. Given that TG concentrations were also less in persistently-hyperglycemic obese dogs, it was hypothesised that TG might play a role in compensatory hyperinsulinaemia (and that a lack of TG could cause a decrease in insulin compensation for hyperglycaemia) (23). In a study involving healthy people without insulin resistance, there were differences between a control group, who responded to a TG infusion with an increase in insulin secretion, and a group with a family history of type 2 DM, who experienced a decrease in insulin secretion, as well as marked hepatic insulin resistance (72). This suggests that insulin responses vary greatly in different physiological states, and this should be considered in future studies

FFA concentrations tend to be greater in insulin-resistant humans, since insulin resistance leads to a decreased insulin-mediated inhibition of lipolysis and, as a result, FFA concentrations tend to increase (51). Furthermore, FFA are thought to be mechanistically involved in the pathophysiology of obesity-induced insulin resistance (51). In studies involving dogs with obesity, FFA concentrations increase and contribute to insulin resistance (36,37,46,69). However, in the present study, FFA and TG concentrations were negatively associated. This was unexpected given that increasing TG concentrations were also associated with greater insulin, glucose and fructosamine concentrations, changes that are evocative of ORMD, and ORMD is commonly associated with increased circulating FFA (51). One possible explanation is that the severity of insulin resistance in the dogs of the present study was mild, or in the early stages, primarily affecting the liver but not peripheral tissues (i.e., insulin is not able to inhibit hepatic glucose production but it is effectively inhibiting lipolysis at the peripheral level). This dissociation between hepatic and peripheral insulin resistance has previously been described (73). An alternative explanation for the decreased FFA concentration, such as enhanced FFA oxidation, seems unlikely given the other changes (increasing insulin, glucose, and fructosamine), which are evocative of ORMD.

The positive association between serum TG and cholesterol has been previously described in overweight dogs and was therefore expected.

Given that some breeds were overrepresented in the overweight category, the risk of misinterpreting an effect of the breed with an effect of overweight was a concern. However, the linear models used were corrected for the effect of the breed, and some effects were evident even in overweight dogs within single breeds. A sex effect was not assessed because, besides for Labrador retrievers, the study design meant that sex was covariate with breed. As a result, correction for the breed effect in the ANOVA and in the linear models would automatically correct for any effect of sex in most cases.

One limitation of the current study was that the BCS is an imperfect measure of body fat mass, and is influenced by many factors, including differences in breed morphologies and fat distribution (60). Further, subjectivity when assigning BCS might have led to the misclassification of some dogs, especially those examined at different centres. However, the same investigator was responsible for the assessment in each one of the five centres, and in all cases they were highly trained veterinarians that are expected to correlate greatly (74). Therefore, the associations found in the present study further support the utility of the BCS as a universal system to estimate fat excess. Different analysis combinations could have been used with BCS categories, but BCS 6 was chosen since it is a widely-accepted cut-off in every-day practice (7,17,18,60,75).

When assigning dogs to the overweight or lean categories, there was a predominance of dogs in the lean category in most breeds (≥80% in most breeds, except for CKCS, NF and LAB, which approached 50% in each category). Such an imbalance might have made it more difficult to identify statistically significant differences between lean and overweight categories within specific breeds that had a small number of overweight dogs. Of course, the limitations for comparisons made between overweight and lean categories would not apply to comparisons with TG concentrations, whose measurement is more accurate than the assessment of BCS. This might have ensured a greater statistical power, explaining the increased number of significant associations with the metabolic variables assessed.

Some may argue that an effect of the diet should have been included in the analyses. However, this study was designed not to test for this parameter, but rather assess a cohort with heterogeneous diets, representative of a client-owned population of dogs. Therefore, no restriction was imposed in terms of diet, as long as the dogs had only access to a commercial dry food diet and received no supplement during the three weeks preceding the analyses.

## Conclusions

Both BCS and serum TG concentrations were independently associated with changes in markers of lipid and glucose metabolism in this large cohort of slightly overweight, otherwise healthy dogs. Non obese, overweight (6-7) body condition was positively associated with insulin and cholesterol concentrations whilst TG concentrations were positively associated with cholesterol, insulin, glucose, and fructosamine concentrations, and negatively associated with FFA concentrations. Therefore, both BCS and fasting TG concentration seem to be useful markers of ORMD-related changes in dogs. Further analyses using more complex multivariate models are needed to better characterize the interplay between these biochemical analytes.

## Acknowledgements

Belgian police and its contribution to the collection of dogs.

## Supporting information

**S1 File. The protocol**.

**S2 File. The dataset**.

